# A cranio-incudo joint as the solution to early birth in marsupials and monotremes

**DOI:** 10.1101/2020.04.08.032458

**Authors:** Neal Anthwal, Jane Fenelon, Stephen D. Johnston, Marilyn B Renfree, Abigail S Tucker

## Abstract

Mammals articulate their jaws using a novel joint between the dentary and squamosal bones. In eutherian mammals, this joint forms in the embryo, supporting feeding and vocalisation from birth. In contrast, marsupials and monotremes exhibit extreme altriciality and are born before the bones of the novel mammalian jaw joint form. These mammals need to rely on other mechanisms to allow them to feed. Here we show that this vital function is carried out by the earlier developing, cartilaginous incus of the middle ear, abutting the cranial base to form a cranio-mandibular articulation. The nature of this articulation varies between monotremes and marsupials, with monotremes retaining a double articulation, similar to that described in the fossil mammaliaform, *Morganucodon*, while marsupials use a versican rich matrix to stabilise the jaw against the cranial base. These findings provide novel insight into the evolution of mammals and the relationship between the jaw and ear.

## Introduction

In non-mammalian vertebrates the craniomandibular (jaw) joint is formed between the quadrate (or palatoquadrate) in the skull and the articular part of Meckel’s cartilage in the mandible. During the evolutionary transition that gave rise to mammals, the connection between the quadrate (the homologue of the mammalian incus) and the cranial base simplified, so that a complex structural attachment of the quadrate to five separate skeletal elements, able to bear the mechanical force of feeding, became a ligamentous suspension of the incus from a single bone, the petrosal, in an air filled cavity allowing sound transmission (Kemp, 2005; Kielan-Jaworowska et al., 2004).

Early mammal-like reptiles had both a permanent Meckel’s cartilage and joints between the quadrate and articular (Q-A), and the quadrate and petrosal in the cranial base – similar to extant reptiles. In mammaliaforms, such as *Morganucodon*, both a Q-A and dentary squamosal joint are present, with a joint between the crista parotica of the petrosal and the incus. The petrosal and incus joint precedes detachment of the middle ear from Meckel’s cartilage in mammal evolution (Luo and Crompton, 1994). A connection between the future middle ear bones and the cranial base is therefore a feature of fossil mammaliaforms. The crista parotica forms as a cartilaginous spur off the otic capsule and is derived from neural crest cells, distinct to the rest of the capsule which is mesodermally derived (O’Gorman, 2005; Thompson et al., 2012). Modern mammals have separated the middle ear from the jaw, as described in (Anthwal et al., 2017; Urban et al., 2017), and the ossicles are now suspended by ligaments from the cranial base to allow free vibration during sound transmission from the ear drum to the inner ear. This is possible due to the novel mammalian jaw joint - the temporo-mandibular (TMJ) or squamosal dentary joint – which provides support between the jaw and cranial base. Paleontological evidence indicates that the evolution of the definitive mammalian middle ear (DMME) occurred at least twice, once in the lineage that gave rise to monotremes and once in the therian (marsupial and eutherian) mammals (Meng et al., 2016; Rich et al., 2005), while new developmental data suggests that the two subclasses of therian mammals may have each independently acquired the DMME (Urban et al., 2017).

Marsupials (Allin, 1975; Filan, 1991) and monotremes (Griffiths, 1978), exhibit extreme altriciality, greater than is seen in any eutherian (Werneburg et al., 2016). This has profound consequences for early feeding as the bones that form the mammalian jaw joint, the dentary and squamosal, have not fully ossified by the time of birth/hatching. The dentary-squamosal joint forms prior to birth in eutherian mammals, and begins to function in the embryo. (Habib et al., 2007; Jahan et al., 2014). In the mouse, gestation is approximately 20 days, with breakdown of Meckel’s cartilage, to isolate the jaw from the ear bones, following during early postnatal stages (Anthwal et al., 2013). In contrast, in the opossum *Monodelphis* the dentary-squamosal is absent at birth, which occurs at around 13 days gestation (Keyte and Smith, 2008), and forms between 14-20 days after birth (Filan, 1991; Maier, 1987). Monotremes hatch out of the egg after 10 days post-oviposition (Griffiths, 1978). The formation of the dentary-squamosal joint in monotremes has recently been followed and shown to form from 10 days after hatching in the platypus (Anthwal and Tucker, 2020).

Given the lack of a jaw joint, it has been proposed that marsupials use the connection between the middle ear bones and cranial base to permit feeding prior to the formation of the secondary jaw joint and cavitation of the middle ear (Crompton and Parker, 1978; Maier, 1987; Sánchez-Villagra et al., 2002).

The feeding strategies of new-born mammals varies in extant members of each subclass of mammals. Compared to eutherian mammals, marsupials rely on placental support for a relatively shorter period of time (Renfree, 2010) and consequently receive the nutrition required for their development via a lengthy and sophisticated lactation (Tyndale-Biscoe and Janssens, 1988; Tyndale-Biscoe and Renfree, 1987). During their early postnatal life marsupials attach to the mother’s teat and use the comparatively early developed tongue musculature to suck (Smith, 1994). In the grey short-tailed opossum, *Monodelphis domestica*, pups are born after 13 days of embryonic development, which is followed by around 14 days permanently attached to the mother’s teat, after which they detach intermittently from the mother but continue to suck. Weaning occurs around postnatal day 60 (Keyte and Smith, 2008).

In contrast to therian mammals, young extant monotremes do not obtain milk in quite the same way as therian mammals due to the absence of teats in the mother (Griffiths, 1978). Instead young monotremes suck up milk vigorously from the flattened but protuberant nipple-like areola on the mother’s abdomen (Griffiths, 1978). In the case of echidnas, these areolae are within the pouch.

Here we describe the cranio-mandibular articulation in monotremes (platypus *Ornithorhyncus anatinus* and short-beaked echidna *Tachyglossus aculeatus*) as they develop from hatching, and compare them to a marsupial (grey short-tailed opossum, *Monodelphis domestica*) and a eutherian (mouse, *Mus musculus*) to ask how these mammals are able to feed prior to the development of the dentary-squamosal joint. Since monotremes and marsupials have different feeding modalities in early life (the sucking from the abdomen vs attached to the mother’s teat), we expect that any role that the middle ear has in the craniomandibular articulation may differ across subclasses. Therefore, we investigate the connection between the malleus and incus, and the incus and cranial base during early postnatal/posthatching development in the echidna, platypus, opossum and mouse.

## Materials and Methods

### Animal tissues

Opossum (*Monodelphis domestica*) tissue was collected as previously described (Anthwal et al., 2017; Urban et al., 2017).

Archival platypus *(Ornithorhynchus anatinus)* and short-beaked echidna *(Tachyglossus aculeatus*) slides were imaged from the collections at the Cambridge University Museum of Zoology, and the Hill Collection, Museum für Naturkunde, Leibniz Institute for Research on Evolution and Biodiversity, Berlin. Details of samples imaged are in Table 1. All museum samples have been studied in previously published works (Green, 1937; Presley and Steel, 1978; Watson, 1916). Stages for platypus are estimated based on Ashwell (Ashwell, 2012). Staging of echidna H.SP EC5 and H.SP EC4 are estimated by cross-referencing (Griffiths, 1978; Rismiller and McKelvey, 2003). Post-hatching day 0 and 3 echidna samples were collected from Prof. Marilyn Renfree, University of Melbourne and Assoc. Prof. Stephen Johnston University of Queensland.

**Table 1:** Museum held specimens used in the current study. CRL – Crown rump length.

\*Estimate based (Ashwell, 2012). <sup>†</sup>Estimate based on (Griffiths, 1978) and (Rismiller and McKelvey, 2003).
| **Species** | **Collection** | **ID** | **Estimated Age** | **CRL / Max length** |
| --- | --- | --- | --- | --- |
| *Ornithorhynchus anatinus* | Cambridge | Specimen W | 2 days * | 16.5mm |
| *Ornithorhynchus anatinus* | Berlin | M45 | 6.5 days * | 33mm |
| *Ornithorhynchus anatinus* | Cambridge | Specimen Delta | 10 days * | 80mm |
| *Ornithorhynchus anatinus* | Berlin | M038 | 30 days | Unknown |
| *Ornithorhynchus anatinus* | Cambridge | HP | 50 days * | 200mm |
| *Ornithorhynchus anatinus* | Cambridge | Specimen Beta | 80 days* | 250mm |
| *Ornithorhynchus anatinus* | Cambridge | HX | 120 days | 295mm |
| *Tachyglossus aculeatus* | Cambridge | Echidna H.SP<br>EC5 | 18 days<sup>†</sup> | 83mm |
| *Tachyglossus aculeatus* | Cambridge | Echidna H.SP<br>EC4 | 55-65 days <sup>†</sup> | 174mm |

Wildtype and *Mesp1Cre; mTmG* were kept at the King’s College London Biological Services Unit. *Sox9CreERT2:tdTomato* embryos were a gift of Prof Robin Lovell-Badge and Dr Karine Rizzoti at the Francis Crick Institute, London.

Phosphotungstic acid (PTA) contrasted embryonic *Pterobnotus quadridens* bat μCT scans were provided by Prof Karen Sears and Dr Alexa Sadier at the University of California Los Angeles.

Guinea pig (*Cavia porcellus*) displays samples were collected as previously described (Anthwal et al., 2015).

Gecko and mouse samples were investigated during embryonic development (35 days post oviposition, 35dpo and E16.5 respectively). The gestation for geckos is around 60 days, and mice have a gestation of 20-21 days. Much of opossum and echidna development occurs during early post-hatching life, including formation of the secondary jaw joint (the TMJ), and so 4 day old opossums and 3 day old echidnas were investigated before the onset of the TMJ.

#### Tissue processing and Histological staining

All tissues for histological sectioning were fixed overnight at 4 °C in 4 % paraformaldehyde (PFA), before being dehydrated through a series of graded ethanol, cleared with Histoclear II, before wax infiltration with paraffin wax at 60°C. Wax embedded samples were microtome sectioned at 8 μm thickness, then mounted in parallel series on charged slides. For histological examination of bone and cartilage, the slides were then stained with picrosirius red and alcian blue trichrome stain using standard techniques.

#### Immunofluorescence

For immunofluorescence staining slides were rehydrated through a graded series of ethanol to PBS. Heat induced antigen retrieval was carried out by microwaving the samples for 10 min in 0.1M Sodium citrate pH6 buffer. Slides were then blocked in 1 % Bovine serum albumin, 0.1 % cold water fish skin gelatine, 0.1 % triton-X for 1 h. Sections were then treated over night at 4 °C with primary antibodies. The following primary antibodies were used, rabbit anti Sox9 (Chemicon) at a dilution of 1/200, chicken anti GFP (Abcam) at a dilution of 1/500, rat anti RFP (Chromotek) at a dilution of 1/200, Rabbit anti Beta-catenin (Santa Cruz) 1/200, mouse anti type 2 collagen (DSHB) at 1/50, mouse anti CD44 (DSHB) at 1/50, mouse anti Tenascin C (DSHB) at 1/40, mouse anti versican (DSHB) at 1/50, rabbit anti versican V1 (Abcam) at 1/400. Following repeated PBS washes, secondary antibodies were added. For fluorescent labelling the following antibodies were used at 1/300: Alexa568 conjugated Donkey anti-Rabbit, Alexa 488 conjugated Donkey anti-Rabbit, Alexa568 conjugated Donkey anti-Mouse, Alexa568 conjugated Donkey anti-Rat, Alexa488 conjugated Donkey anti-Chicken (all Invitrogen). Secondary antibodies were added in the blocking buffer for 1 h at room temperature in the dark. The secondary antibody was then washed off with PBS, and the slides mounted with Fluroshield mounting medium containing DAPI (Abcam). Sections were visualised by Leica SP5 confocal microscopy. For Versican and CD44 slides, secondary biotinylated goat anti-mouse antibody (Dako) was added to the slides 1/400 in blocking buffer. Slides were then washed in PBS before being treated with ABC-HRP streptavidin kit (Vector Labs), and then revealed with DAB (Vector Labs). Monotreme immunofluorescence staining was carried out in technical replicates due to the rare nature of the samples. Mouse and opossum analysis was carried out in biological triplicates.

#### In situ hybridisation

Radioactively labelled antisense RNA probes were made against mouse *Gdf5* and *Bapx1* mRNA, and radioactive in situ hybridisations were carried out to detect the expression of these genes in sagittal plain cut sections of wildtype mice, as previously described (Tucker et al., 2004). All in situ staining was carried out in biological replicates.

#### 3D Reconstruction

Three-dimensional reconstructions of middle ear and surrounding cranial base cartilages were generated from serial histology images in FIJI (ImageJ 1.47v), using the Trackem2 Plugin (Schindelin et al., 2015, 2012).

#### Cell density counts

Cell density was counted in a DAPI stained sections of 5 day old opossums (n=3). In FIJI, 20 separate 80μm^2^ fields were randomly placed across the mesenchyme surrounding the incus across 5 sections. The total number of nuclei were counted if they were located wholly within the field, or where more than 50% of the nuclei intersected the upper or right hand margin of the field. Next by looking at parallel alcian blue stained sections, the fields were scored as being in proteoglycan rich or weak regions. Two fields were ambiguous, and so were removed from the analysis. The user did not know the proteoglycan status of the field at the time of counting. Next the mean cell number in each field was calculated in the remaining 8 proteoglycan rich (alcian blue stained) regions and 10 proteoglycan weak (weak alcian blue stain) regions, and compared by unpaired two tailed students t-test in Prism statistical analysis software (Graphpad).

## Results

### The primary jaw joint (malleus-incus) does not provide a site of articulation in marsupials and monotremes at birth

In mice (*Mus musculus*), the malleus and incus are initially formed from a single cartilaginous condensation that separates, by the formation of a joint, at Embryonic (E) day 15.5 (Amin and Tucker, 2006). At birth, therefore, the incus and malleus are evident as distinct cartilages (Figure 1B). In *Monodelphis domestica*, the malleus and incus are still connected at birth at the dorsal end by a ridge of cartilage (Filan, 1991) (Figure 1C). We observed a similar connection between the malleus and incus in the echidna (*Tachyglossus aculeatus*) just after birth. Like the opossum, the middle ear ossicles were fused dorsally, indicating that they function as a unit (Figure 1D). The homologous elements in reptiles form a clear synovial joint in the embryo, as shown in the ocelot gecko (*Paroedura picta*) (Figure 1A). These findings demonstrate that, like opossums, monotremes do not use the primary jaw joint as the craniomandibular articulation before the development of the dentary-squamosal joint

**Figure 1.**
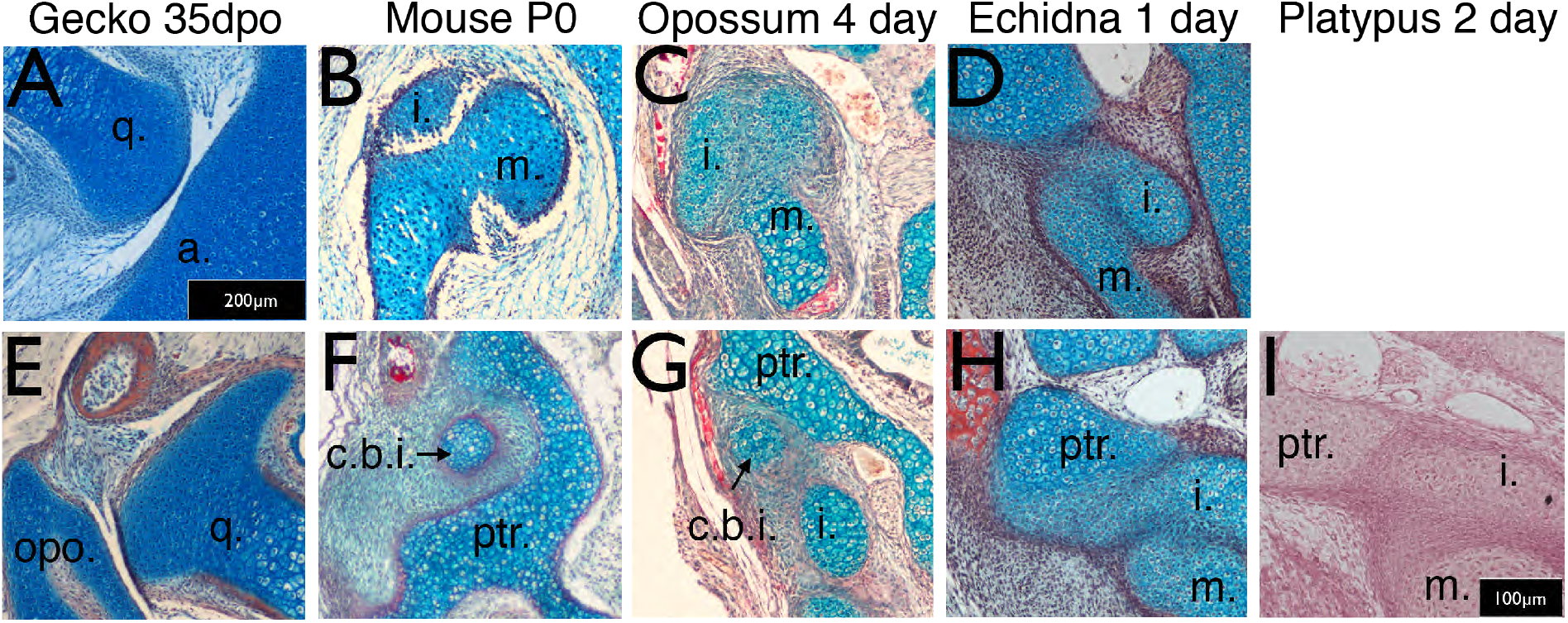
Timing of the develop of the quadrate-articular / malleus-incus, and cranio-incudo joints. Histological sections stained with alician blue and picrosirius red. The primarily jaw articulation is formed by 35 days of post-oviposition (35dpo) during in ovo development in geckos (A). The malleus-incus joint, the homologue of the quadrate-articular joint, is formed during in utero development in mice, and is fully formed at birth (Po) (B). The malleus incus joint is still partially fused in 4 day postnatal opossum pups (C) and 3 day post-hatching echidna young (D). During development the gecko quadrate forms a joint with the opisthotic (E). At birth there is no articulation between the crus breve of the incus in mice (F}, not is an articulation observed in P4 opossums (G). The incus is fused with the petrosal in both P3 echidna (H) and the P2 platypus. q. quadrate; a. articular; i: incus;. m. malleus; opo, opithotic; c.b.i crus breve of the incus; ptr. petrosal.

In marsupials, the true joint prior to the formation of the dentary-squamosal joint appears to be between the incus and the cranial base (Maier, 1987). We therefore investigated the relationship between the incus and the petrosal in the cranial base in mice, opossums, platypus and echidna, comparing the interaction to the developing joint between the quadrate and opisthotic in embryonic geckos. In many reptiles, as shown in the gecko, the quadrate (incus homologue) forms a synovial joint with the opisthotic (also known as the otoccipital) in the cranial base during embryonic development (Figure 1E). The opisthotic/otoccipital is morphologically equivalent to the petrosal of mammals. In mice, the crus breve (short process) of the incus nestled in a fossa created by the crista parotica of the petrosal, but was separated by a region of mesenchymal cells, highlighting the lack of a clear articulation point between the two elements (Figure 1F). The incus at birth, therefore only physically contacted the adjacent middle ear bones, the malleus and stapes. Similar to the mouse, the crus breve in neonatal opossums, fitted into a fossa created by the crista parotica, but abutted the petrosal on the inferior aspect of the crista parotica (Figure 1G). The incus and petrosal were therefore positioned much closer than in the mouse.

The relationship between the incus and crista parotica in the two monotreme species was significantly different from the other mammals. In both platypus (*Ornithorhynchus anatinus*) and echidna (*Tachyglossus aculeatus*), the incus appeared to be fused with the crista parotica at birth (Figure 1H, I). The lower jaw, via Meckel’s cartilage, would therefore be physically connected to the upper jaw, via the incus at this timepoint. 3D reconstructions of the incus, malleus and petrosal, showing the relationship of these different elements in the different species is shown in Supplementary Figure 1. The relatively small size of the incus in both monotremes is striking, as is the extended and tapered crus breve of the incus in the opossum.

**Figure 1 Supplementary.**
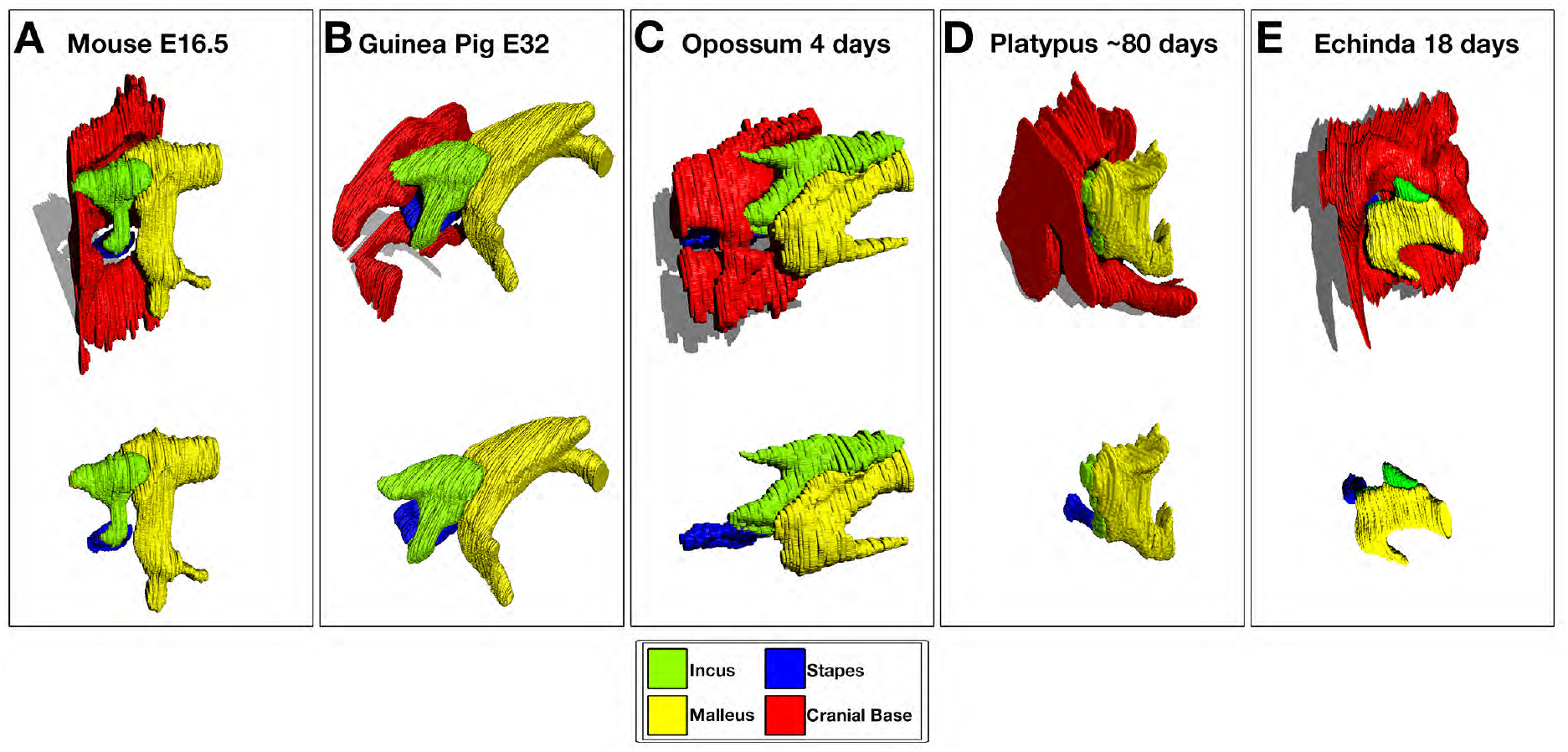
3D Reconstruction of cartilaginous middle ear ossicles and cranial base from histological sections shows differences in anatomy in different orders of mammals during development.

### Development of an incus-petrosal joint in monotremes during early feeding

To investigate the monotreme relationship between the incus and crista parotica further we followed development of these two cartilages from birth to functional use of the dentary-squamosal joint, but before cavitation of the middle ear space. At 2 days and 6.5 days the platypus incus was fused to the crista parotica by immature chondrocytes (Figure 2A,B). Between 10 days and 30 days the connection was difficult to make out, with the two cartilages almost completely integrated together (Figure 2C, D). Strikingly, by 80 days, when the dentary-squamosal joint would have started to be functional, the incus and crista parotica were no longer fused, with the two distinct cartilages abutting each other (Figure 2E). At this stage, in contrast to the other stages investigated, the ear ossicles and petrosal had begun to ossify. However, the regions forming the malleus-incus joint, and the incus-petrosal articulation remained as cartilage. A cartilaginous articular surface between the incus and petrosal was maintained at 120 days, a period when the young would have started to leave the burrow (Figure 2F)(Holland and Jackson, 2002). A similar move from early fusion, to articulation was observed in the echidna (Figure 2F-J). No evidence of a synovial capsule, however, was identified at any stage.

**Figure 2.**
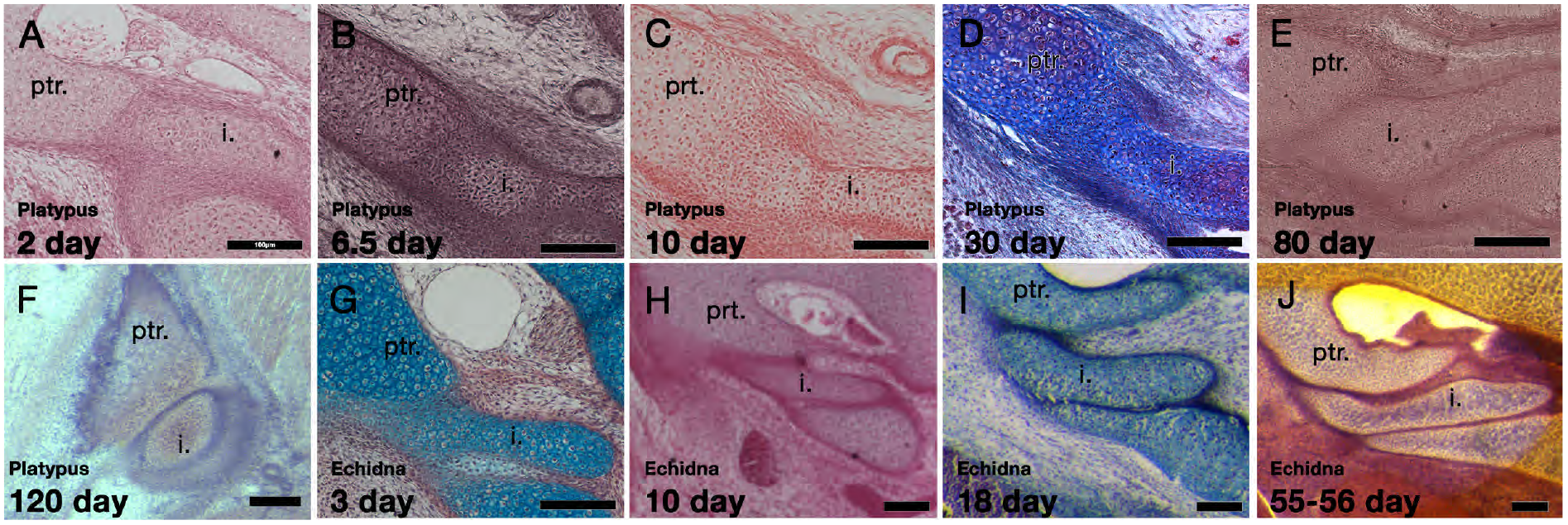
Development of the incus-petrosal joint in monotremes. The platypus incus is fused to the petrosal by immature chondrocytes at 2 days (A) and 6.5 days (B). At 10 days, the fusion persists, with mature chondrocytes forming the connection (C). A similar morphology is seen at P30 (D). At 80 days the incus and petrosal are no longer fused, but instead the two cartilages abut each other (E). At 120 days the incus and petrosal have begun to ossify, but the region of articulation in between the two elements remains cartilaginous (F). In echidna the incus is fused to the petrosal by immature chondrocytes at 3 days (G) and 10 days (H). By 18 days the two elements are separated but remains abutted (I). This connection remains though to 55-65 days(J)

The fusion of the incus and crista parotica coincides with the period when the young would have been feeding from milk, while the move to an articulation was associated with periods when the dentary-squamosal is fully formed and functional (Figure 2 supplementary). This suggests a period where two cranial-mandible connections would have been functional in the platypus - between Meckel’s cartilage and the petrosal, and between the dentary and squamosal.

**Figure 2 Supplementary.**
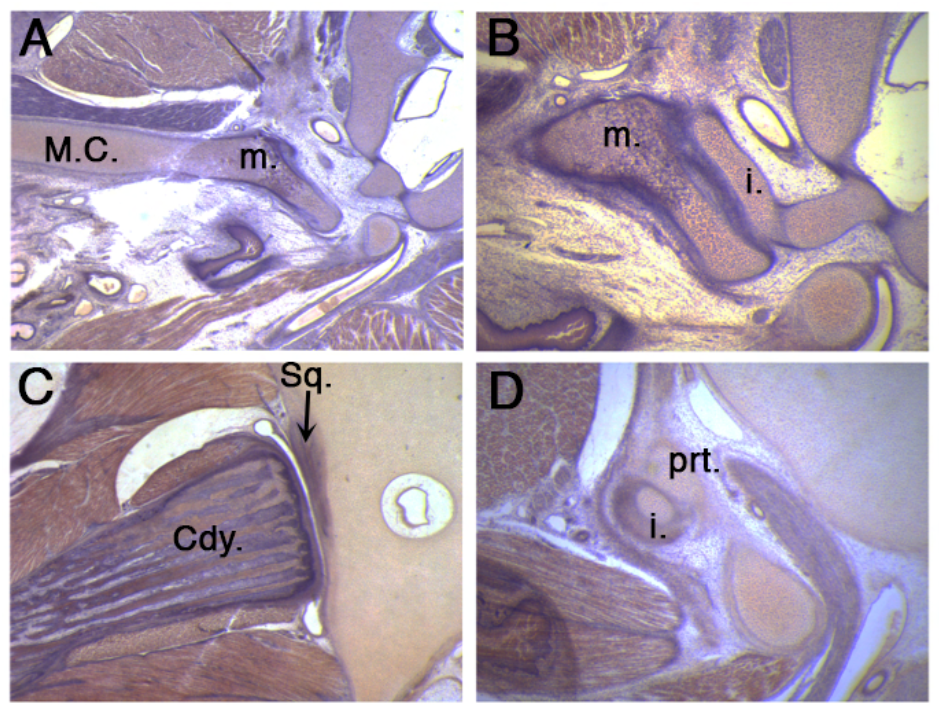
50 day platypus showing an intact connection between Meckel’s cartilage and the malleus (A), a un-cavitated middle ear (B), a fully formed synovial jaw joint (C) and an articulation between the incus and petrosal in the cranial base (D). M.C. Meckel’s cartilage, m. malleus; i. incus; Cdy. Condylar process of the mandible bone; Sq. squamosal bone; ptr, petrosal.

Middle ear cavitation occurred very late in the monotreme specimens analysed, with only the 120 day platypus showing partial cavitation around the hypotympanum, but this did not extend up to the attic where the ossicles are housed. Hearing, thus, must be a very late developing sense in the platypus.

### Upregulation of Wnt signalling initiates joint formation between the ossicles and cranial base in echidna

In order to further understand the change in the relationship between the incus and petrosal, immunohistochemistry staining was carried out in 0 day old and 3 days old echidna samples.

In the fused incus-petrosal region of 0 day old echidna (Figure 3A), the expression of both a master regulator of cartilage development, Sox9, and a principal component of cartilage extra cellular matrix, Collagen Type 2, were continuous between the incus and the crista parotica of the petrosal, as well as between the incus and the malleus (Figure 3B). Since the connection between these elements is lost later in post-hatching development, IF for beta-catenin was carried-out. Nuclear localised beta-catenin is a readout of canonical Wnt signalling, and is known to negatively regulate chondrocytes differentiation and promote joint formation (Hartmann and Tabin, 2001)Few beta-catenin positive cells were observed within the cartilage of the middle ear and petrosal at 0 days, though beta-catenin was strongly expressed in the neuro-epithelium of the inner ear (Figure 3C). At post-hatching day 3, the incus and crista parotica were still fused, although the cells joining the two elements resembled fibrocartilage or immature chondrocytes (Figure 3D). Expression of Sox9 was still strong and continuous throughout all elements (Figure 3E, E’), however Collagen Type 2 expression was weaker in the fusion region (Figure 3E, E”), possibly indicating a change in cartilage type from hyaline cartilage to fibrocartilage. Interestingly nuclear beta-catenin, suggestive of active Wnt signalling, was observed in two strips, in the chondrocytes between the incus and petrosal, and within the malleus-incus joint, indicating suppression of cartilage fate in these regions (Figure 3F). Upregulation of Wnt signalling between the incus and petrosal therefore, may play a role in formation of a joint between these two, initially fused, structures.

**Figure 3.**
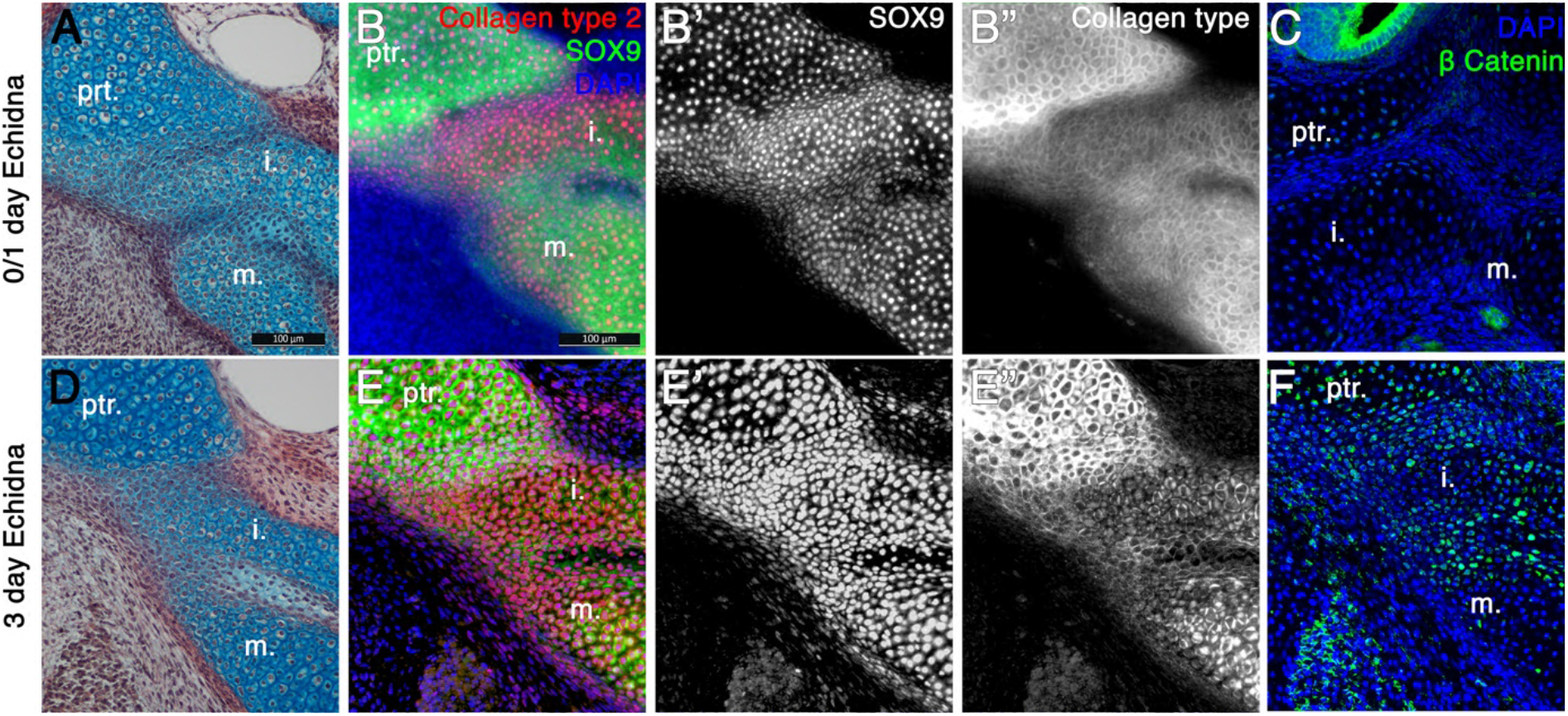
Fusion of the Incus with the petrosal in Echidna pouch young. A: alcian blue / picrosirius red staining on the fusion between the incus and petrosal observed in the newly hatched echidna. B: Immunohistochemical staining against the regulator of chondrogenesis Sox9 (green) (B,B’) and the marker of mature cartilage Collagen type 2 (red) (B,B”) demonstrates that the cartilaginous incus and pertrosal bones are fully fused at post-hatching day 0. C: Immunohistochemistry again the β Catenin (green) shows no activity within the cartilages at this timepoint. Expression is observed in the neuroepithelium of the inner ear. D: alcian blue / picrosirius red staining on the fusion between the incus and petrosal observed in 3 day old echidna shows that the elements are now fused by fibrocartilage. E: Immunohistochemical staining against the regulator of chondrogenesis Sox9 and the marker of mature cartilage collagen type 2 (E,E”). Sox9 is still continuously expressed between the elements (E,E’), but collagen type 2 is down regulated in the incuspetrosal and incusmalleus articulation region (E,E”). F: Immunohistochemistry again the β Catenin shows nuclear localisation within the incus-petrosal and incus-malleus articulation regions, indication active canonical Wnt signalling, an important step in suppression of chondrogenesis during joint formation. ptr. petrosal; i. incus; m. malleus.

### Interactions between the petrosal and incus are also observed prenatally in eutherian mammals

While the fusion between the incus and petrosal in echidna and platypus could be explained by the evolutionary distance between monotremes and therian mammals, it has also been suggested that the incus is transiently attached to the cranial base in 7 week old human fetuses (Rodríguez-Vázquez et al., 2018).. This suggests that the potential for fusion may be a default state in mammals. In order to examine this, we next undertook fate mapping experiments in the mouse, and investigated the relationship between the incus and petrosal in other eutherian mammals during embryonic development.

Potentially chondrogenic Sox9 expressing cells were fate mapped by tamoxifen induction at E14.5 in *Sox9CreERT2; tdTomato* mice, which were then collected at P0. At this stage Sox9 (green) was expressed in the petrosal and incus and suspensory ligaments, overlapping with the red fluorescent Protein (RFP) marking the Sox9 lineage cells. In addition, the red Sox9 lineage cells were found in the Sox9 negative mesenchymal cells, in the gap between the petrosal and incus (Figure 4A). A pre-cartilaginous bridge is therefore evident in the mouse between the incus and the crista parotica. Next, expression of Sox9 was investigated at E14.5. The incus, and the crista parotica are both neural crest derived (O’Gorman, 2005; Thompson et al., 2012), while the rest of the petrosal is mesodermal. We therefore looked at the expression of Sox9 (red) in *Mesp1Cre; mTmG* mice, where mesoderm derived tissue expresses GFP (Figure 4B). Sox9 protein was expressed continuously between the incus and the petrosal. The incus Sox9 expression domain was continuous with the expression domain of the neural crest derived crista parotica, which in turn is fused to the mesodermal portion of the petrosal. Since the incus does not fuse with the petrosal in the mouse, despite the expression of Sox9 between the elements, we next looked at the mRNA expression of joint markers *Gdf5* and *Bapx1* between the incus and petrosal of mice by in situ hybridisation (Figure 4C-E). *Gdf5* was expressed in the mesenchyme between the incus and petrosal, as well as in the malleus-incus joint (Figure 4D). *Bapx1*, which specifies both the malleus-incus joint and the quadrate-articular joint (Tucker et al., 2004), was not expressed in between the incus and the petrosal (Figure 4E). In the mouse, therefore there is a potential for the incus and crista parotica to fuse but they are prevented from doing so by the upregulation of the joint marker Gdf5.

**Figure 4.**
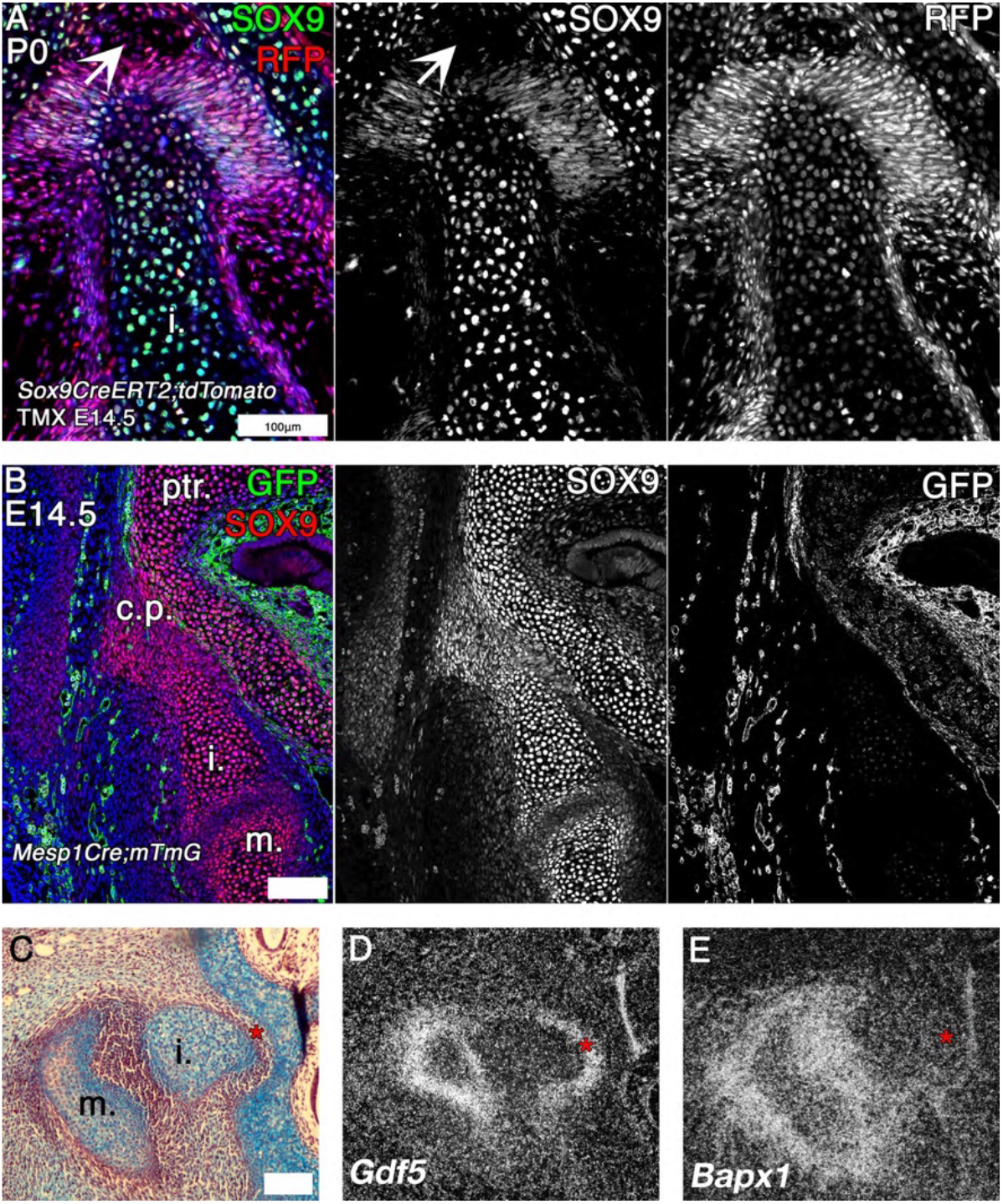
Mouse fate mapping studies demonstrate developmental fusion between incus and petrosal. A: Genetic tracing of chondrogenic Sox9 expression cells by inducible reporter mice at postnatal day 0 (P0). Sox9 lineage cells (red) are observed in the mesenchyme and developing ligaments between the crus breve of the incus and the petrosal. Sox9 protein (green) is not expressed in the mesenchyme surrounding the incus at P0 (arrow head). B: Genetic tracing of mesoderm lineage cells (green) and immunohistochemistry against Sox9 protein (red) at embryonic day 14.5 (E14.5). Sox9 expression at E14.5 confirms that the incus and petrosal are formed of a continuous chondrogenic mesenchyme, and that the incus joins with the petrosal at the crista parotica, which is not of mesodermal origin. C-E Expression by in situ hybridisation of joint markers in sagittal section of E14.5 mouse middle ears. *Gdf5* mRNA is expressed with the malleus-incus joint, and between the incus and the petrosal (D), potentially acting to inhibit the fusion maturation of the Sox9 expressing mesenchyme between the ear the cranial base into cartilage. The middle ear joint marker *Bapx1* is not expressed between the incus and the petrosal (E). * indicates space between of incus and petrosal in C-E. ptr. petrosal; i. incus; m. malleus.

Very close associations between the incus and crista parotica during development were also observed in other eutherian mammals via PTA stained microCT (see bat in Figure 4 Supplementary), suggesting that interactions between these two elements are observed as a feature prenatally in eutherian mammals, similar to post-hatching monotremes. The function of this prenatal connection between the upper and lower jaw is unclear but may act as a brace to buffer movement during this period.

**Figure 4 Supplementary.**
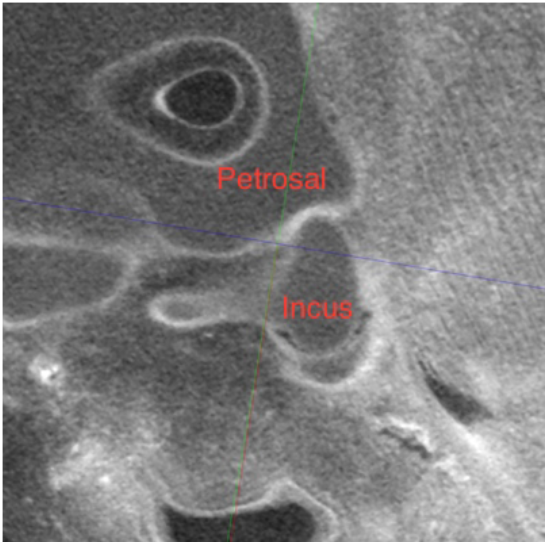
Contrast enhanced μCT of embryonic bat (*Mormoops blainvillei*) middle ear at Carnegie Stage21, showing abutment of the crus breve of the incus against the petrosal

### Petrosal-incus relationships in marsupials

Next we investigated the articulation between the incus and petrosal observed in the developing opossum. It was originally suggested that the marsupial incus forms a joint with the crista parotica (Maier, 1987), although this was disputed in *Monodelphis* (Filan, 1991). Although this later paper found no evidence of a joint they did show the mesenchyme between the crista parotica and incus as being condensed (Filan, 1991). We therefore, investigated the extra cellular matrix (ECM) components of the mesenchyme surrounding the opossum incus in more detail. It was noted that mesenchyme surrounding the crus breve and superior portion of the body of the incus had a more intense staining with alcian blue compared to those regions round the inferior border of the incus and the other ossicles (Figure 1C, 2C,G). This pattern was observed throughout ossicle development (5A-C). In order to further characterise the differences in the ECM in the different regions of the middle ear mesenchyme, immunohistochemistry for versican was carried out. Versican is a large proteoglycan with side chains of glycosaminoglycans (GAGs), such as hyaluronic acid (HA). Proteoglycan complexes act to attract water, and are held in place by collagen fibres to stiffen the matrix in hyaline cartilage, and act to lubricate articular cartilage (Wu et al., 2005). Versican is required during the initial condensation of mesenchyme but is absent from mature cartilage, where aggrecan is expressed (Kamiya et al., 2006). Veriscan expression is maintained in the joint region during limb cartilage development, acting to inhibit maturation of the mesenchyme to form cartilage (Choocheep et al., 2010; Snow et al., 2005)

Versican was strongly expressed in the mesenchyme surrounding the short arm of the incus at 5 days, 10day and 27days, correlating with the region of strong alcian blue expression (Figure 5D-F). The high level of versican around the crus breve, therefore suggests a role for the ECM in providing a buffering function in this region.

**Figure 5.**
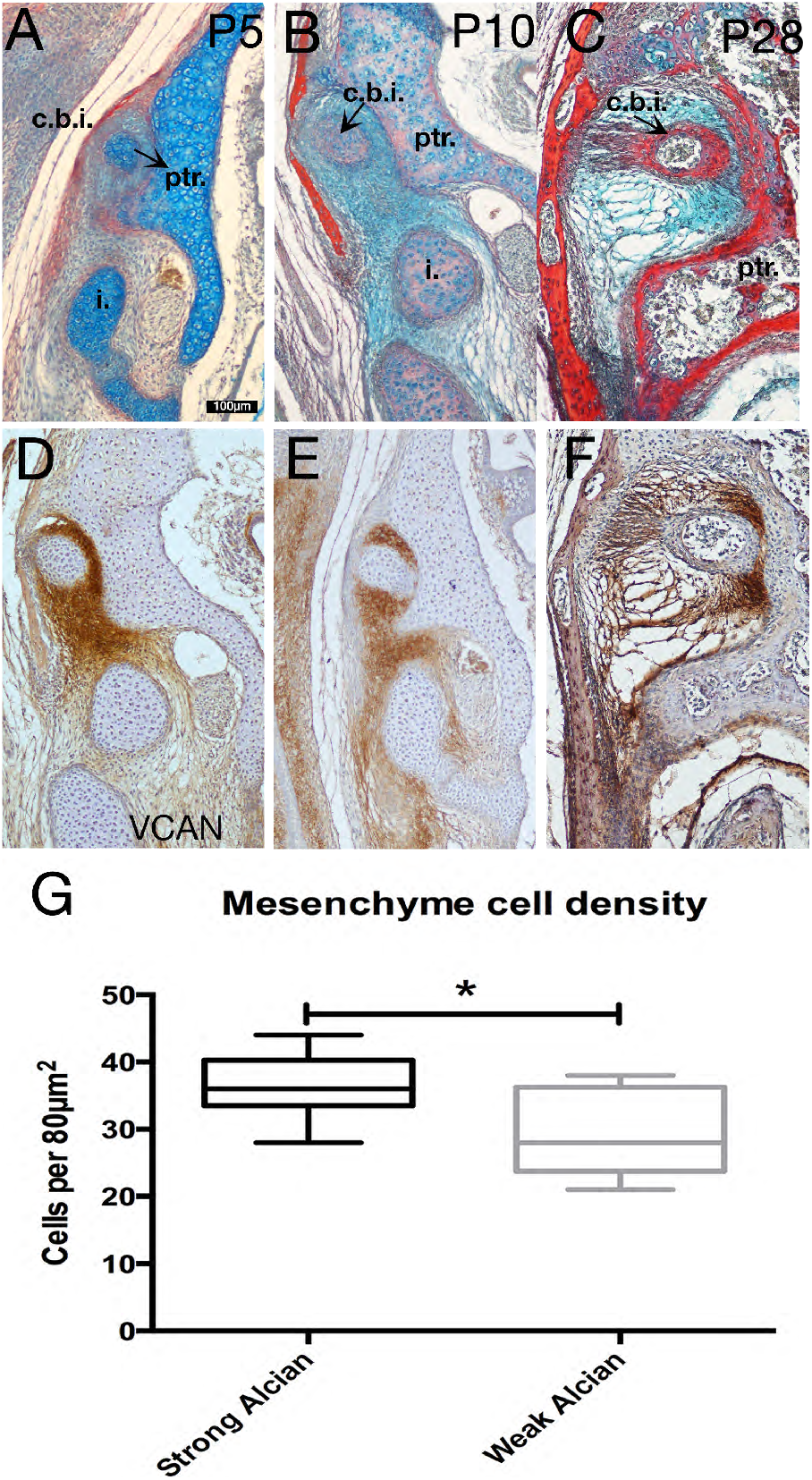
Specialist mesenchyme supports incus-petrosal connection in juvenile opossums. A-F Mesenchyme surrounding the crus breve of the incus is rich in the proteogclycan Versican (Vcan) at 5 days (A,D) and 10 days (B,E). During cavitation of the middle ear versican rich mesenchyme is concentrated between the crus breve of the incus and the petrosal (C,F). G: At day 5 the proteoglycan rich regions surrounding the crus breve have a significantly greater cell density than the regions with less proteoglycan.* P=0.0152 unpaired two tailed t-test. Error bars = 1 SD. i. incus; c.b.i crus breve of the incus; ptr. petrosal

Cell density of the mesenchyme was measured in regions with strong alcian blue /versican staining and compared against the cell density of regions with low alcian blue / versican staining. Unpaired two tallied t-test demonstrates that the regions with high alcian blue had a significantly higher (P=0.0152) cell density than those regions with lower alcian staining (figure 5G).

**Figure 5 Supplementary.**
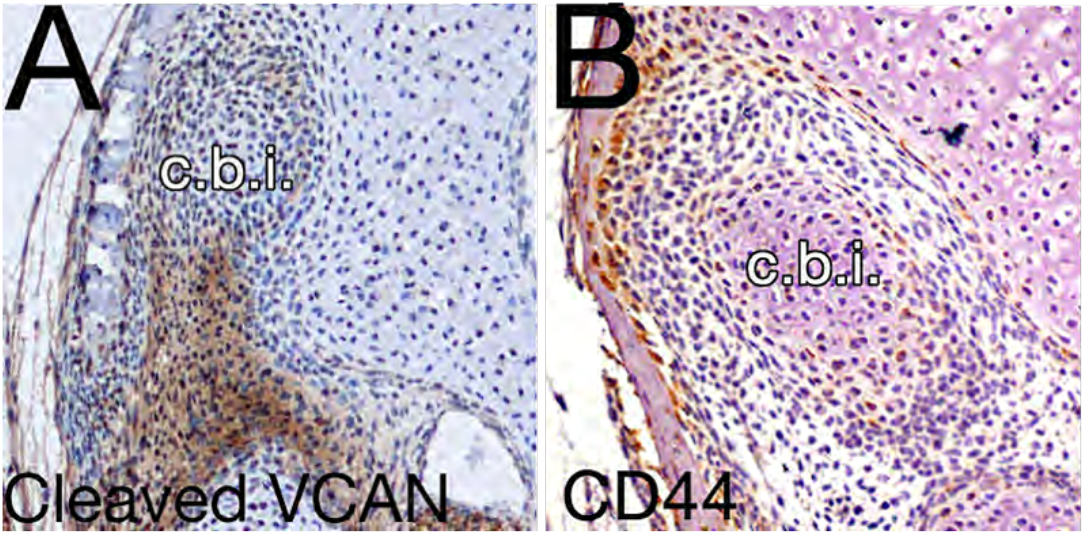
Expression by immunohistochemistry of cleaved veriscan DPEAAE (A) and CD44 (B) in 10 day opossums at the level of the crus breve of the incus. c.b.i. Crus Breve of the Incus.

Versican is processed by ADAMTS family members for clearing and remodelling (Nandadasa et al., 2014). While the full length form of versican is thought to have a structural role, the cleaved form has a an active role in signalling, influencing morphogenesis and tissue remodelling (Nandadasa et al., 2014). Interestingly when we analysed the cleaved form of versican, using antibodies against DPEAAE, the expression was largely reciprocal to that of uncleaved versican, with lower levels specifically around the crus breve (Supplementary Figure 5A). This suggests that versican around the incus is protected from cleavage allowing it to maintain its structural role. The lack of cleaved versican around the crus breve, suggests the lack of a signalling role in this region, in agreement with the low level of expression of CD44, a cell surface receptor and binding partner of versican-hyaluronan complexes. CD44 was not associated with the mesenchyme around the crus breve, but was instead restricted to the perichondrium of the cartilaginous elements and periosteum of the skeletal elements of the ear (Supplementary Figure 5B).

## Discussion

We demonstrate here that the crus breve of the incus in monotremes is fused to the petrosal during early post hatching life (Figure 2, Suppl. Figure 2, Figure 3), and that the joint marker nuclear beta catenin expression correlates with the separation of these elements during the early post-hatching stages (Figure 3F). Fate mapping and gene expression studies in mice indicate that the crus breve of the incus and the crista parotica are formed from a continuous region of Sox9 expressing chondrogenic cells (Figure 4 A, B). *Gdf5*, an early joint marker (Storm and Kingsley, 1999) and regulator of chondrogenesis (Francis-West et al., 1999), was expressed in the region between the crus breve and crista parotica in the mouse (Figure 4C). Furthermore, the incus and cranial base temporarily fuse during the development of the human middle ear region (Rodríguez-Vázquez et al., 2018). Together these data indicate that the fusion of the incus to the cranial base is not a derived feature of the monotremes, and that the common mammal-like reptile ancestors of both monotremes and therian mammals may have formed an articulation between the quadrate/incus and petrosal though fusion of the elements followed by joint formation though Wnt and Gdf5 signalling.

The current study indicates that the first pharyngeal arch derived incus forms a continuous field of chondrocytes with the second arch derived crista parotica, which in turn is fused with the mesoderm derived body of the petrosal. The borders between these developmentally distinct populations are, therefore, not always reflected by the mature anatomy.

### Marsupial opossum has a specialised anatomy to brace middle ear against cranium during sucking

The crus brev of the incus is elongated in the developing opossum compared with other species (Supplementary Figure 1). In order to feed by suckling in the absence of a TMJ we propose that this anatomy allows for an increased surface contact with the cranial base during postnatal development, which, in combination with the proteoglycan rich surrounding mesenchyme, acts to stabilise the mandible against the rest of the head. It is noted that many adult marsupials have a relatively elongated crus breve of the incus compared to eutherian species, for example the bare-tailed woolly opossum *Caluromys philander,* and the grey short tailed opossum, *Monodelphis domestica* (Sánchez-Villagra et al., 2002). Even when eutherian mammals have a longer crus breve, such as in Talpid moles, the process is thinner and more finger-like compared to that of marsupials (Segall, 1973, 1970). This may be a consequence of the developmental requirement for an elongated short process to facilitate feeding before the development of the mature mammalian jaw articulation.

In the majority of adult marsupials, including *Monodelphis*, the incus is suspended from the cranial base by the suspensory ligaments, and the crus breve extends into a fossa. One interesting exception is the marsupial mole, the crus breve of which has a connective tissue attachment to a lamella on the petrosal (Archer, 1976). This results in the middle ear ossicles being affixed to the cranial base, an adaptation to a fossorial niche found in other mammals such as in true moles. In light of the current study, the absence of an incudal fossa in the marsupial mole may be interpreted as a retention of the juvenile petrosal morphology (paedomorphy).

### Consequence of ECM in opossum middle ear

In adult non-mammalian amniotes the homologue of the incus -the quadrate- and cranial base are strongly attached by fibrous syndesmoses or cartilaginous synchondroses (Payne et al., 2011), and we show that a synovial joint appears to form in geckos during development (Figure 1). In the neonatal opossum neither type of connection is observed. In neonatal marsupials Sánchez-Villagra and colleagues describe the connection between the incus and petrosal as being an “immature syndesmosis”, which acts as a “supportive strut” during sucking (Sánchez-Villagra et al., 2002). In the current study we demonstrate a specialised condensed mesenchyme surrounds the incus of opossum postnatal juveniles. We show that this condensed mesenchyme is rich in the proteoglycan versican (Figure 5). In contrast expression studies in human foetuses demonstrate that versican is restricted to the perichondrium of Meckel’s cartilage (Shibata et al., 2014, 2013), with high hyaluronic acid levels within the joints but not surrounding the incus (Takanashi et al., 2013). This concentration of versican around the crus breve therefore appears to be a feature of *Monodelphis*, and perhaps marsupials in general.

The versican-rich mesenchyme may act to either stabilise the incus by increasing the tension of the surrounding mesenchyme during feeding, “lubricate” the articulation between the incus and cranial base by increasing the hydration of the ECM, or both. In keeping with this role, versican is dynamically expressed at the pubic symphysis during pregnancy in mice (Rosa et al., 2012), during which time the mouse pubic symphysis forms a fibrous joint or syndesmosis (Ortega et al., 2003). Significantly, there is little cleaved versican (DPEAAE) around the crus breve of the incus, suggesting a mechanical, rather than a signalling role (Figure 5 Supplementary A). Overall it is likely that this mesenchyme is supporting the incus, rather than enabling mobilisation, with the high level of uncleaved versican acting to increase fibroviscocity while also elevating hydration of the ECM. In this way, the mesenchyme around the incus acts as a cushion during the mechanical stress of suckling.

For young monotremes and marsupials, the middle ear must function as part of the mandible until the dentary-squamosal bones have formed. This is similar, but not homologous to the situation in cynodont ancestors of mammals. In these animals, the quadrate/incus articulated with a number of cranial elements, including the quadratojugal, to stabilise the jaw articulation. These connections and many elements like the quadratojugal have been lost in extant mammals in order to free the incus and increase its mobility during sound transmission. The mechanical requirements for feeding placed upon the middle ears in monotremes and marsupials during early life have resulted in the fusion of the incus and petrosal in monotremes, and the elongated contact supported by a proteoglycan matrix in marsupials. These adaptations allow for stabilisation of the middle ear before the development of the TMJ and separation of the middle ear from the mandible, but do not compromise the effectiveness of the middle ear in later life.

### A double jaw articulation during monotreme development

We have demonstrated here that juvenile monotremes have two connections between the mandible and cranial base. The first connection is through the middle ear, which in juveniles remains attached to the mandible and is fused with the cranial base via the incus. The second is the later developing novel mammalian jaw joint-the TMJ. Only much later in the life of the young does it appear that the connection between the middle ear and mandible is lost, and the malleus and incus act as a DMME. This novel finding has significant implications for the evolution of the middle ear and jaw joint in mammals. Fossil evidence indicates that mammalian ancestors had a persistent connection between the middle ear ossicles and the jaw, as evidenced by the presence of an ossified Meckel’s element, or a dentary groove and post dentary trough, supporting a persistent Meckel’s cartilage (Luo, 2011; Rich et al., 2005; Urban et al., 2017). For these animals, the connection of the middle ear with the jaw took one of two forms, in each case the mammalian secondary jaw joint was present. The first was a more basal mandibular middle ear where the incus and malleus were firmly attached to the cranial base and dentary respectively. More derived fossils had a partial, or transitional mammalian middle ear (PMME or TMME), where the middle ear was medially inflected away from the dentary, presumably allowing for improved vibration, but the malleus was still connected to the jaw, via Meckel’s cartilage (Luo, 2011). In these fossils with a PMME, little is understood of the rear of the ossicular chain, where the incus meets the petrosal, due to the poor and rare preservation of middle ear ossicles in the fossil record, a consequence of their small size. For example, only recently has a multituberculate with a complete incus been described (Wang et al., 2019). Our data suggests that even in these transitional mammals with a PMME, the incus would have still articulated with the cranial base via the crista parotica, at least at some point during the animal’s life history.

The DMME appears to have evolved independently in monotremes and therian mammals (Rich et al., 2005). Due to the absence of evidence we do not know if the incus articulation in animals with a PMME varied in a lineage specific manner, with the therian lineage resembling juvenile marsupials, and monotremaformes resembling juvenile platypuses and echidna, or if both lineages had similar articulation. The data from transgenic reporter mice (Figure 4), along with data from humans (Rodríguez-Vázquez et al., 2018) and non-model therians (Figure 4 supplementary) suggests that the monotreme-type fusion and articulation of the incus with the cranial base may have been common in mammal like-reptiles. If so, the developing monotreme, with a double jaw articulation and a fused or articulated incus and petrosal, provides an exciting model for the study of the developmental basis of mammalian evolution.

## Acknowledgements

Thanks to Karen Sears (UCLA) for the bat microCT scan. Thanks to Robert Asher for access to monotreme samples held at the Zoological Museum in Cambridge University, and to Andrew Gilis for imaging. Thanks to Peter Giere for access to the Hill Collection at the Berlin Museum fur Naturkunde. Thanks to Prof Robin Lovell-Badge and Dr Karine Rizzoti at the Francis Crick Institute, London for provision of the Sox9cre/Tom mice.

## Author contributions

NA and AST conceived and planned the work. NA and AST were involved in acquisition of the data. JF, SDD, MBR were involved in provision of the freshly fixed echidna samples. All authors played a role in interpretation of the data. AS and NA drafted the paper, with input from AJF and MBR.

